# Dynamic Modulation of Beta-Band Oscillations in the LGN and Their Role in Visual Processing

**DOI:** 10.1101/2025.07.08.663741

**Authors:** Henry J. Alitto, Alyssa N. Sanchez, Prescot C. Alexander, W. Martin Usrey

**Affiliations:** Center for Neuroscience, University of California, Davis, Davis, CA, 95618, USA; Department of Neurobiology, Physiology, and Behavior, University of California, Davis, Davis, CA, 95618, USA; Helen Wills Neuroscience Institute, University of California, Berkeley, Berkeley, CA, 94720; Department of Neuroscience, University of California, Berkeley, Berkeley, CA, 94720; Information Systems and Modeling Group (A-1), Los Alamos National Laboratory, Los Alamos, NM, 87545, USA

**Keywords:** thalamus, cortex, contrast gain control, arousal, spatial attention

## Abstract

Neuronal oscillations are a ubiquitous feature of thalamocortical networks and can be dynamically modulated across processing states, enabling thalamocortical communication to flexibly adapt to varying environmental and behavioral demands. The lateral geniculate nucleus (LGN), like all thalamic nuclei, engages in reciprocal synaptic interactions with the cortex, relaying retinal information to and receiving feedback input from primary visual cortex (V1). While retinal excitation is the primary driver of LGN activity, retinal synapses represent a minority of the total synaptic input onto LGN neurons, allowing for both retinogeniculate and geniculocortical signals to be influenced by nonretinal sources. To gain a holistic view of network processing in the geniculocortical pathway, we performed simultaneous extracellular recordings from the LGN and V1 of behaving macaque monkeys, measuring local field potentials (LFPs) and spiking activity. These recordings revealed prominent beta-band oscillations coherent between the LGN and V1 that influenced spike timing in the LGN and were statistically consistent with a feedforward process from the LGN to V1. These thalamocortical oscillations were suppressed by visual stimulation, spatial attention, and behavioral arousal, strongly suggesting that these oscillations are not a feature of active visual processing. Instead, they appear analogous to occipital lobe, alpha oscillations recorded in humans and may represent a signature of signal suppression that occurs during periods of low engagement or active distractor suppression.

**Significance Statement:** Oscillations within thalamocortical networks in the awake state are generally believed to enhance communication between the thalamus and cortex, allowing circuits to flexibly respond to changes in sensory, behavioral, and cognitive demands. Here, we show that oscillations within and between the LGN and V1 are suppressed by increases in visual stimulation, increases in behavioral arousal, and shifts in covert spatial attention. We therefore conclude that these oscillations are not a mechanism to enhance the transmission of retinal information through the LGN to V1. Instead, we propose that they are a signature of signal suppression that occurs when network engagement is low or during active distractor suppression.

## Introduction

The thalamus is the inextricable partner of the cortex, serving as a critical hub for sensory and cognitive information flow from the periphery to the cortex, as well as providing a transthalamic pathway for communication between cortical areas (Jones, 1985; Sherman and Guillery, 2013; Usrey and Sherman, 2023). Indeed, all thalamic relay nuclei have reciprocal connections with the cortex, and all cortical areas have reciprocal connections with the thalamus. Neuronal oscillations are a ubiquitous feature of thalamic network activity, varying with sensory, behavioral, and cognitive demands (Bazhenov et al., 2000; Steriade, 2006; Saalmann et al., 2012; Zhou et al., 2016; Bazinet et al., 2021; Jayachandran et al., 2023). For higher order thalamus, including the pulvinar, oscillations are thought to modulate thalamocortical communication, allowing networks to adapt to changing demands (Saalmann et al., 2012; Sweeney-Reed et al., 2014; Helfrich and Knight, 2016; Halassa and Kastner, 2017; Hwang and Shine, 2023). Whether oscillations serve similar roles in first-order thalamus remains unclear.

In the visual system, the lateral geniculate nucleus (LGN) of the dorsal thalamus is a first order nucleus and the primary source of retinal information to the cortex, receiving monosynaptic excitation from the retina and sending monosynaptic excitation to primary visual cortex (V1; Usrey and Alitto, 2015). While the LGN’s primary role is relaying retinal signals to V1, this transmission is not absolute; instead, retinal signals are selectively gated according to visual processing demands and behavioral state (Usrey et al., 1998; O’Connor et al., 2002; McAlonan et al., 2008; Rathbun et al., 2010; Alitto et al., 2011, 2019; Wimmer et al., 2015; Fisher et al., 2017; Crombie et al., 2024). This gating is influenced by non-retinal inputs, which comprise the majority of LGN synapses, including cortical feedback from V1, GABAergic inhibition from local interneurons and the thalamic reticular nucleus, and cholinergic input from the brainstem (Harting and Guillery, 1976; Steriade et al., 1988; Guillery and Harting, 2003; Wang et al., 2006; Usrey and Alitto, 2015; Krueger and Disney, 2019).

Previous work identified coherent beta-band oscillations between the LGN and V1 in behaving macaques (Bastos et al., 2014). While these geniculocortical oscillations exhibit temporal features consistent with feedforward transmission, it remains unknown whether they are modulated by network demands. To address this, we quantified how geniculocortical oscillation magnitude changes as the LGN becomes more engaged in visual processing due to increased visual stimulation or behavioral demands. We hypothesize that if these oscillations actively facilitate the transmission of retinal information through the LGN to V1, then they should increase in magnitude as the strength of the visual stimulus increases and the behavioral significance of the visual stimulus increases.

To gain a holistic understanding of oscillations in the geniculocortical pathway, we recorded neuronal activity from behaving macaques and monitored local in the LGN and V1 across behavioral and visual conditions. Consistent with previous reports (Bastos et al., 2014), we found robust beta-band oscillations within the LGN that were coherent with oscillations in retinotopically neighboring locations in V1 and were consistent with a feedforward thalamocortical process. We further found that these beta-band oscillations influenced spike timing within the LGN, providing a mechanism for these oscillations to be communicated from the LGN to V1. In contrast to the hypothesis that these oscillations facilitate visual processing, we observed that they were suppressed during periods of increased LGN processing demands. In particular, the geniculocortical oscillations were suppressed by increased visual stimulation, heightened arousal, and shifts of spatial attention toward the LGN receptive field center. These dynamics resemble those of occipital alpha oscillations in human EEG, which may represent a signature of signal suppression that occurs when processing demands are low or during active distractor suppression. A possible source for these oscillations is the thalamic reticular nucleus, which provides GABAergic inhibition to the LGN and is modulated by behavioral arousal and attention (Guillery and Harting, 2003; McAlonan et al., 2008; Wimmer et al., 2015).

## MATERIALS AND METHODS

Six adult rhesus monkeys (*Macaca mulatta*; four female and two male) were used for this study. All experimental procedures conformed to National Institutes of Health and United States Department of Agriculture guidelines and were approved by the Institutional Animal Care and Use Committee at the University of California, Davis. Under full surgical anesthesia, the animals received a cranial implant consisting of a head post for head stabilization and two recording cylinders that allowed access to the LGN and V1.

### Electrophysiological recordings

Extracellular recordings from the LGN and V1 were made with platinum-glass electrodes (1–2 MV; Alpha Omega) using a microdrive (40 mm electrode travel; Thomas Motorized Electrode Manipulator, Thomas RECORDING) mounted on the recording chamber. Continuous voltage signals were amplified (A-M Systems), filtered (0.1–5 kHz), and recorded using a Micro1401 data acquisition system (28 kHz) and Spike2 software (Cambridge Electronic Design, CED). To extract local-field potentials (LFPs), extracellular recordings were low-pass filtered (stopband edge frequency = 250 Hz) with a 6th degree, Chebyshev filter (Matlab: cheby2, filtfilt). LFP signals were subsequently subsampled to 500 Hz and notch filtered at 60 Hz to remove mains noise (Matlab, custom software).

### Visual stimulation and receptive field mapping

Visual stimuli were generated with a ViSaGe (Cambridge Research Systems) and presented on a gamma-corrected CRT monitor (Sony or Mitsubishi) positioned ~80 cm in front of the animal. The display had a resolution of 1024×768, a refresh rate of 100-140Hz, and a mean luminance of ~38 cd/m2. Receptive fields of LGN and V1 neurons were manually mapped using small (<0.5°), user-controlled, computer-generated visual stimuli (custom software with Spike2 interface).

### Behavioral training and performance

Data was collected while the animals performed either a passive fixation or a spatial attention task. For both tasks, eye position and pupil size were monitored using an Eye-Trac6 infrared, eye tracker (Applied Science Laboratories) with a custom Spike2 interface. For passive fixation, the animals fixated on a small (~0.2° diameter) fixation dot, using a fixation window of 0.8°-1.2° diameter. During passive fixation, sine wave gratings varying in stimulus contrast, spatial frequency, and size were presented centered over the retinotopic coordinates of the recorded neuron’s receptive field.

### Spatial attention

Two animals were trained to perform a spatial attention task. For this task, the animals initiated a trial by fixating a central fixation dot similar to the fixation dot used for passive fixation. When the animal obtained fixation, two sine wave gratings (temporal frequency = 5 Hz, spatial frequency = 1.0 cycles/°) appeared at equidistant locations from the fovea, typically at an eccentricity of 4-8°. Each sine wave grating was surrounded by a colored circle (one green, one red) and the color of the circle at each location was constant throughout the task. The color of the fixation point (either red or green) indicated to the animal which grating was 95% likely to change contrast (valid trials) after a delay with an exponential hazard function of 0.6 – 1.8s. On 5% of the trials (invalid trials), the uncued grating increased in contrast (same hazard function). The base contrast of the grating was set to be near the C50 (contrast to evoke a half-maximum response) of the recorded LGN neuron, as estimated through hand mapping, and the contrast change was titrated so that the animal performed ~ 85% correctly. After the contrast change, the animal was given 1s to respond by making a saccade to the location of the change, and the animal was rewarded for correct responses with a small juice reward. Psychophysical performance (accuracy and reaction time) for valid and invalid trials was compared to ensure that the animals were applying spatial attention as anticipated.

### Analysis

#### Oscillatory LFP activity

LFP power spectra from LGN and V1 recordings were computed using a Fourier transform with a Hanning window (custom Matlab script). To identify significant oscillatory activity, we first estimated the mean and standard deviation of the non-oscillatory, fractal component via a bootstrap analysis of the LFP power spectra. Specifically, we iteratively sampled the power spectrum with replacement 1000 times, drawing the same number of trials as in the original dataset for each iteration, and fitting each to a linear, log-log relationship between power and frequency:

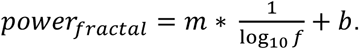

Here, m and b were optimized to capture the linear relationship between log(power) and log(frequency), respectively. From this, the residual oscillatory power was defined as

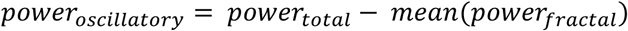

Similarly, the Z-score of the oscillatory power was defined as

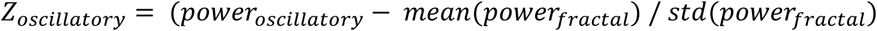

Z-scores greater than 5.0 were taken as evidence of statistically significant oscillatory activity at a given frequency.

#### LGN-V1 phase synchrony

Frequency-specific LGN-V1 field-field phase synchrony was estimated using pairwise phase consistency (ppc, Vinck et al., 2010):

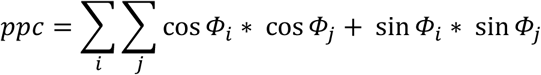

Here, Φ_i_ and Φ_j_ are the phase angles between the LGN and V1 on trials i and j, respectively. Low ppc values near 0 indicate no consistent relationship between LGN and V1 LFP phase, while ppc values near 1.0 indicate a constant phase relationship between LGN and V1.

To determine the significance of PPC values, we estimated the expected standard deviation of PPC under the null hypothesis of no consistent phase relationship. We modeled the null as a random walk, where each step is taken from the distribution of *cos*(*ϕ_i_* − *ϕ_j_*) where *ϕ_i_* and *ϕ_j_* are independently drawn from a uniform distribution over [−π, π]. In this case, *cos*(*ϕ_i_* − *ϕ_j_*) follows an arcsine distribution on [−1, 1], which has a variance of 0.5. If we assume that the expected step size is 0 (Vinck et al., 2010) and the variance of the step size is 0.5, then the total displacement after N steps:

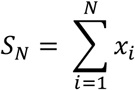

This is a sum of independent and identically distributed variables, so the variance of the total displacement is:

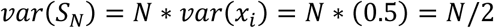

Since PPC considers normalized displacement,

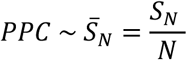

This is the mean of N zero-mean random steps. Thus,

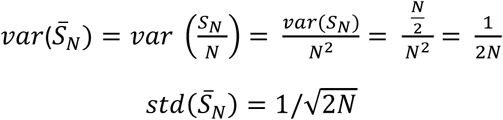

Here, N is the number of pairwise comparisons. PPC values were then z-scored based on this analytically defined N-dependent standard deviation. The validity of this estimation of the null distribution was supported by simulation of PPC values based on random phases using a range of N values (Figure S1). Values exceeding 5.0 standard deviations were considered evidence of significant field-field phase synchrony.

#### Directionality of LGN-V1 communication

To determine if LGN-V1 phase synchrony resulted from either a feedforward or feedback process, we calculated the directionality of LGN-V1 LFP interactions using phase slope index (psi, (Nolte et al., 2008b). Positive and negative psi values are statistically consistent with feedforward and feedback processes, respectively. The significance of psi values was not estimated for individual recordings but instead was estimated for the sample distribution as a whole using a standard t-test to determine statistically significant differences from 0.

#### Spike-field phase synchrony

LGN spike-field phase synchrony was estimated with ppc, using the phase of 0.4s LFP snippets centered on spike times. Because recordings were made using single, high-impedance electrodes, the LFP data was contaminated with residual spike waveforms that could not be removed by lowpass filtering. These spike waveform artifacts potentially interfered with the ability to detect spike-field phase synchrony, particularly in the beta band, where we saw the best evidence for significant oscillatory activity in the LGN. To deal with this issue, we reasoned that the strength of frequency-specific spike-field synchrony would depend upon the presence of strong oscillatory activity in the LGN, while the influence of the spike artifact would be independent of the underlying oscillatory activity. Therefore, we calculated ppc values for epochs of strong and weak beta band oscillatory activity and defined the uncontaminated estimate of LGN spike-field phase synchrony as the difference between these two values.

#### Matching Pursuit

To visualize spike-related transients in the local field potential, we used matching pursuit (mp), a greedy algorithm that decomposes signals into linear combinations of time–frequency localized basis functions (“atoms”) drawn from an overcomplete dictionary. This approach adaptively selects atoms that best match the signal structure, allowing high-resolution representations of both rhythmic and transient features. We implemented the mp algorithm in Matlab following the procedure described by (Chandran et al., 2016), using Gabor atoms spanning a range of scales. Matching pursuit was used solely for visualization and was not involved in any quantitative analyses of spike–LFP phase relationships.

## RESULTS

The lateral geniculate nucleus (LGN) is the primary source of retinal information to the cerebral cortex, receiving monosynaptic excitation from retinal ganglion cells and sending monosynaptic excitation to primary visual cortex (V1). The primary goal of this study was to gain a holistic view of network dynamics in the LGN by recording local field potentials (LFPs, n = 1,117) from behaving macaque monkeys (n = 6) under various behavioral conditions and visual stimulation paradigms. Single-unit spiking activity was also recorded. In a subset of recordings, we simultaneously recorded from both the LGN and V1 at similar retinotopic locations (n = 227). Unless otherwise stated, recordings were made while the monkeys passively fixated on a central location with visual stimuli centered over the LGN receptive field.

As will be shown below, our results from the LGN recordings show a prominent oscillation in the beta-frequency band (low beta = 11-23 Hz, high beta = 24 – 35 Hz), that influenced spike timing within the LGN, was coherent with oscillations in V1, and was statistically consistent with a feedforward, thalamocortical process from the LGN to V1. These thalamocortical oscillations were suppressed by visual stimulation, behavioral arousal, and spatial attention directed toward the LGN receptive field.

### Beta-band oscillations in the LGN

Our first objective was to measure LGN power spectra with the goal of determining whether there are significant neuronal oscillations in the local network. Towards this goal, extracellular recordings were made from six passively fixating macaque monkeys, totaling 888 recordings from both parvocellular and magnocellular layers (Figure 1, column 1). To determine the presence of significant neuronal oscillations, each power spectrum was fit to a 1/f equation on a log-log scale (Figure 1, column 2). This allowed us to estimate and remove the non-oscillatory, fractal component (Figure 1, column 3), leaving an estimate of the oscillatory power. A bootstrap analysis was used to estimate the significance of the residual oscillatory power at each frequency (Figure 1, column 4).

**Figure 1:**
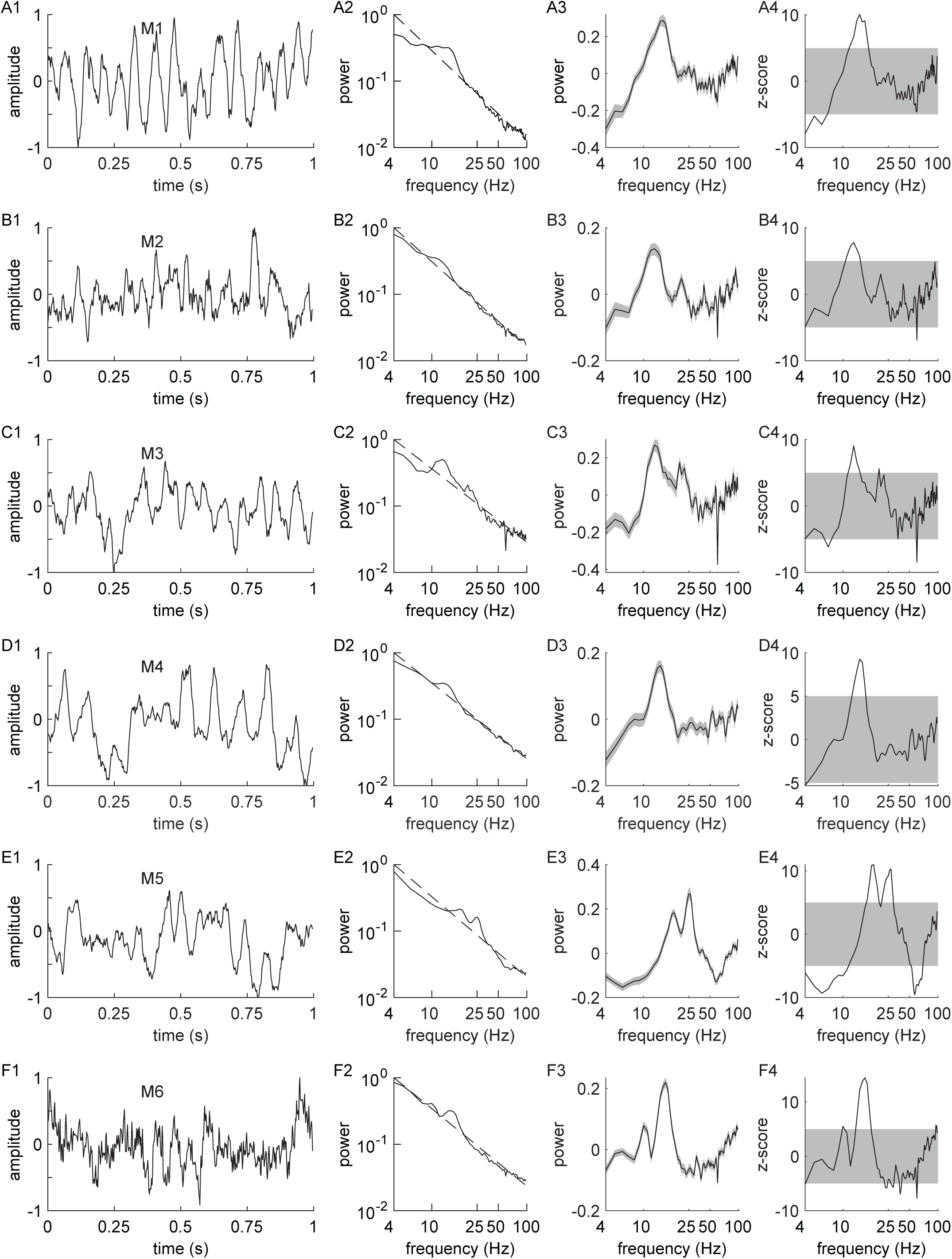
LGN local field potential. (A1) Snippet of an LGN extracellular recording, lowpass filtered (<250 Hz) to extract LFP activity (M1 = animal ID). (A2) Power spectrum from the full, representative recording. Dashed gray line = 1/f fit. (A3) Oscillatory power, defined as the residual power after subtracting the 1/f fit. (A4) Z-score of the oscillatory power. Power above the gray shaded box was considered evidence of significant oscillatory activity (B-F) Additional examples, one from each monkey included in this study.

Neuronal oscillations were most commonly observed in the LGN LFP in the range of 11-35 Hz (Figure 2A and B). This roughly corresponds to the lower (11-23 Hz) and higher (24-35 Hz) segments of the beta-frequency band (Kopell et al., 2000; Jensen et al., 2005; Cohen et al., 2007). Indeed, the distribution of significant frequencies has two clear peaks, one centered over the low-beta range and one centered over the high-beta range (Figure 2B). Across our six animals, 81% of recordings showed significant beta-band oscillations in either the low- or high-beta band (75.9% low-beta, 30.4% high-beta), with a range of 58.8% - 96.3% across animals. In 5 of 6 animals, low-beta oscillations were the dominant neural oscillation (Figure 2A-F, H), with one animal having a near-even distribution of low-beta and high-beta oscillations (Figure 2G). In general, we detected few functional differences between low- and high-beta oscillations. In the sections below, we identify any significant differences between low-beta and high-beta oscillatory properties, using the term beta oscillation when no differences were observed. Less commonly, significant neural oscillations were also seen in other frequency bands. In particular, we observed significant alpha-band oscillations (6-10 Hz), gamma-band oscillations (40-70 Hz), and broad-band, high-frequency oscillations (>70 Hz) in a minority of recordings (17.9%, 10%, 23%, respectively). These oscillations will not be emphasized here but may be of interest for future studies.

**Figure 2:**
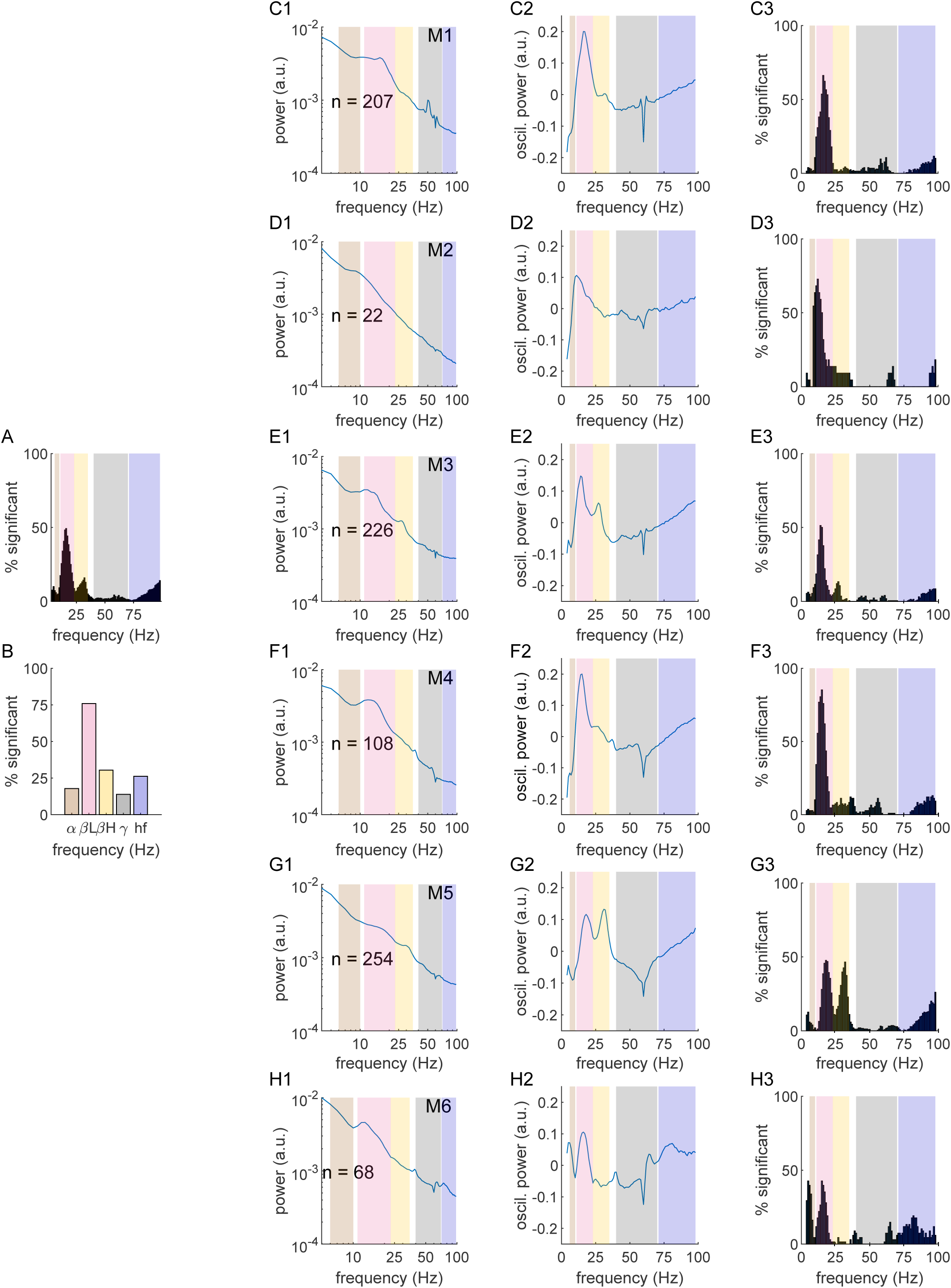
Oscillatory activity in LGN LFP. (A) Distribution of significant residual oscillatory activity in the LGN across all 6 animals. (B). Percentage of LGN recordings with significant oscillatory activity in 5 frequency bands: alpha 7-10 Hz, low-beta 11-23 Hz, high-beta 24-35 Hz, gamma 40-70 Hz, and high-frequency >70 Hz. (C1) Average LGN power spectrum for monkey M1 (n = # of recordings). (C2) Average LGN oscillatory activity for monkey M1. (C3). Percentage of recordings with significant oscillatory activity at each frequency for monkey M1. (D-H) Same as C, one row for each animal included in this study.

### Coherent thalamocortical beta-band oscillations

Given that the primary role of the LGN is to relay retinal information to V1, we next sought to understand the relationship between beta-band oscillations in the LGN and V1. Towards this goal, in a subset of our recordings, simultaneous recordings were made from regions in the LGN and V1 that had a maximum retinotopic displacement of 3.0° visual degrees (minimum = 0.1°, median = 1.4°). From these recordings, pairwise phase consistency (ppc) was used to calculate field-field phase synchrony between the LGN and V1 (see Materials and Methods).

Similar to the distribution of significant neural oscillations within the LGN, significant LGN-V1 field-field synchrony was detected in the beta-band frequency range, as can be clearly seen in the four example LGN-V1 pairs (Figure 3A-D) and the sample average (Figure 3E1). Across all four monkeys, 80.6% and 63.9% of recordings had significant LGN-V1 phase synchrony in the low-beta and high-beta bands, respectively. As expected, the probability of finding significant beta-band LGN-V1 phase synchrony was dependent upon the presence of significant beta-band oscillatory power in the LGN (Z_ßlow_ = 2.17, p = 0.0147; Z_ßhigh_ = 4.9, p < 10^−5^). Similar relationships were seen for the alpha band (Z_alpha_ = 7.0, p < 10^−5^) and the high-frequency bands (Z_alpha_ = 4.6, p = 10^−5^), but not the gamma band (Z_gamma_ = 0.034, p = 0.4866). Further, the magnitude of LGN-V1 beta-band phase synchrony was highly dependent upon the strength of the underlying LGN oscillation (Figure 3E2-3), being significantly stronger during periods of high oscillatory strength compared to periods of low oscillatory strength for both low-beta (ppc_weak_ = 0.10±0.01, ppc_strong_ = 0.17±0.02, p < 10^−5^) and high-beta oscillations (ppc_weak_ = 0.11±0.01, ppc_strong_ = 0.20±0.02, p < 10^−5^) (Figure 3E2-3). This indicates that beta-band coherence is not simply a reflection of recording from two synaptically connected areas. Instead, the dependence of the LGN-V1 phase synchrony on the strength of the underlying LGN oscillation indicates stronger communication between the LGN and V1 when beta oscillations are present. Finally, using phase slope index (psi) as a measure of the directionality of LGN-V1 communication, we determined that beta oscillations were statistically consistent with a feedforward, thalamocortical process from the LGN to V1 (Figure 3F).

**Figure 3:**
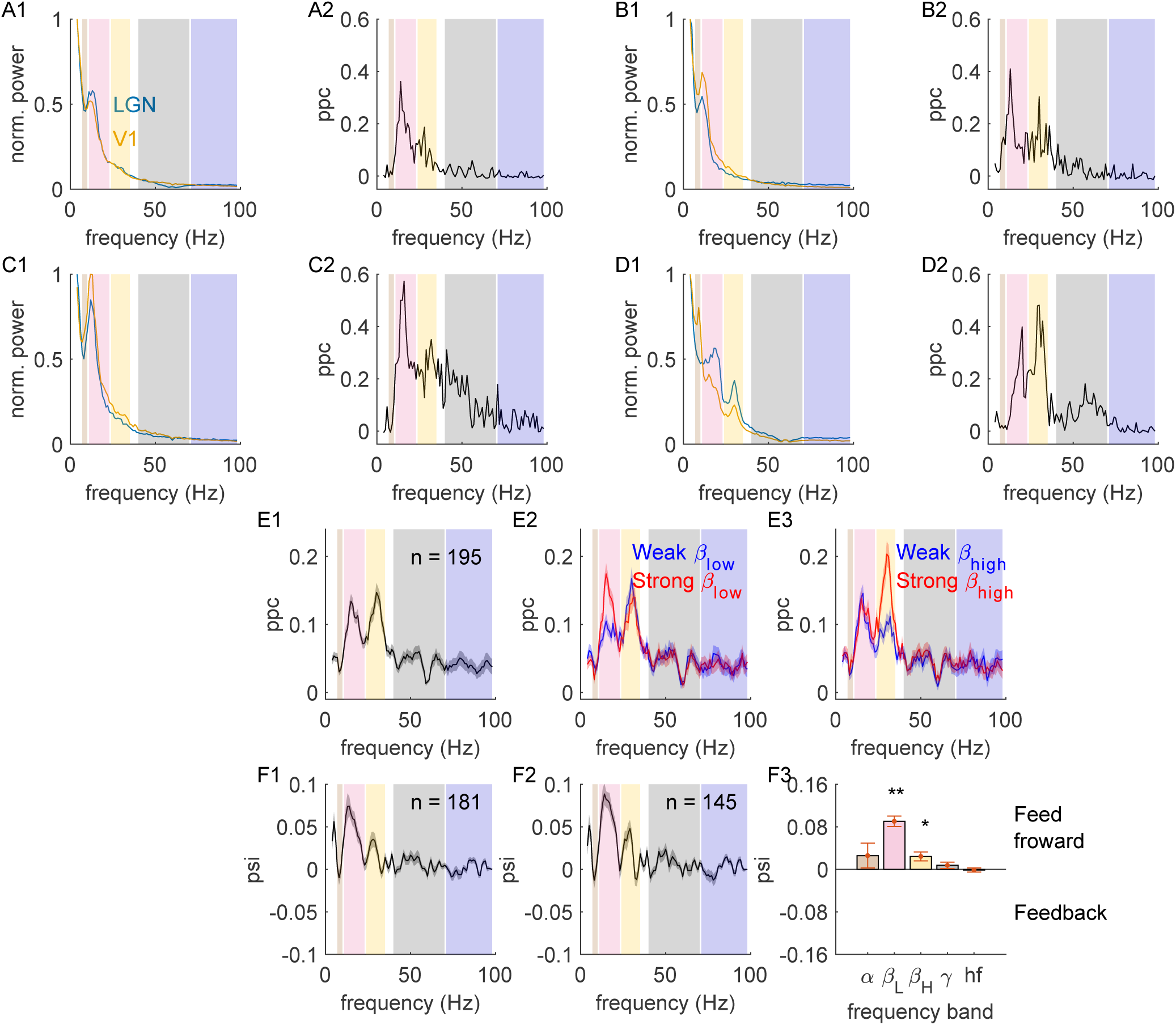
LGN-V1 field-field phase synchrony. (A1) Example of simultaneously recorded LGN and V1 power spectra (LGN = blue, V1 = orange). (A2) Example field-field ppc for paired recording shown in A. (B-D) Additional examples, same as A. (E1) Average LGN-V1 field-field ppc across our sample. (E2) PPC separated into trials with weak or strong beta oscillations (blue trace and red trace, respectively) for recordings with significant low beta-band LGN-V1phase synchrony. (E3) Same as E2, but for recordings with significant high beta-band LGN-V1 phase synchrony. (F1) Average LGN-V1phase slope index (psi) for recording with significant low beta-band LGN-V1 phase synchrony. Positive and negative values are indicative of feedforward (LGN→V1) and feedback processes (LGN←V1), respectively. (F2) Same as F1, but for recordings with significant high beta-band LGN-V1 phase synchrony. (F3) Sample distribution for frequency band specific psi values (* = p< 0.001, ** = p<10^−5^).

### Spike-field phase synchrony in the LGN

We next sought to determine the functional influence of LGN beta-band oscillations on LGN spiking activity. Before describing these interactions, it is useful to consider the involvement of artifactual contributions that can affect the analyses performed and a means to avoid them. The data analyzed for this study were collected with single, high-impedance electrodes, typically with well-isolated single units with large signal-to-noise ratios (Figure 4, column 1). Previous studies have demonstrated that when the spiking activity and LFP are recorded from the same electrode, the residual spike waveform, which cannot be completely removed by high-pass filtering, will cause artifactual interactions between the LFP and spiking activity (Ray et al., 2008; Ray, 2015). Indeed, the influence of residual spike waveforms is evident in the spike-triggered average LFP waveform (Figure 4, column 2), and the residual spike waveform underlies an artificial, broad-band, high-frequency spike-field interaction that dominates the average LGN ppc curves (Figure 4, column 3). The relationship between the spike waveform and the high-frequency coherence can be further visualized using matching pursuit (see Materials and Methods), a spectral analytic technique that allows one to achieve temporal resolution that exceeds the theoretical limits of traditional Fourier analysis. As can be seen from the average matching pursuit across the sample of LGN recordings (Figure 4C), the broadband, high-frequency, spike-field synchrony is restricted to a narrow temporal window (~8ms) surrounding the time of LGN action potentials. This is more consistent with a spike artifact than an ongoing oscillation. Moreover, we found that the strength of the broadband, high-frequency, spike-field phase synchrony was highly correlated with the signal-to-noise of the underlying spike waveform (Figure 4D, r = 0.56, p < 10^−5^). Thus, the residual spike waveform would interfere with the ability to measure spike-field interactions using traditional approaches.

**Figure 4:**
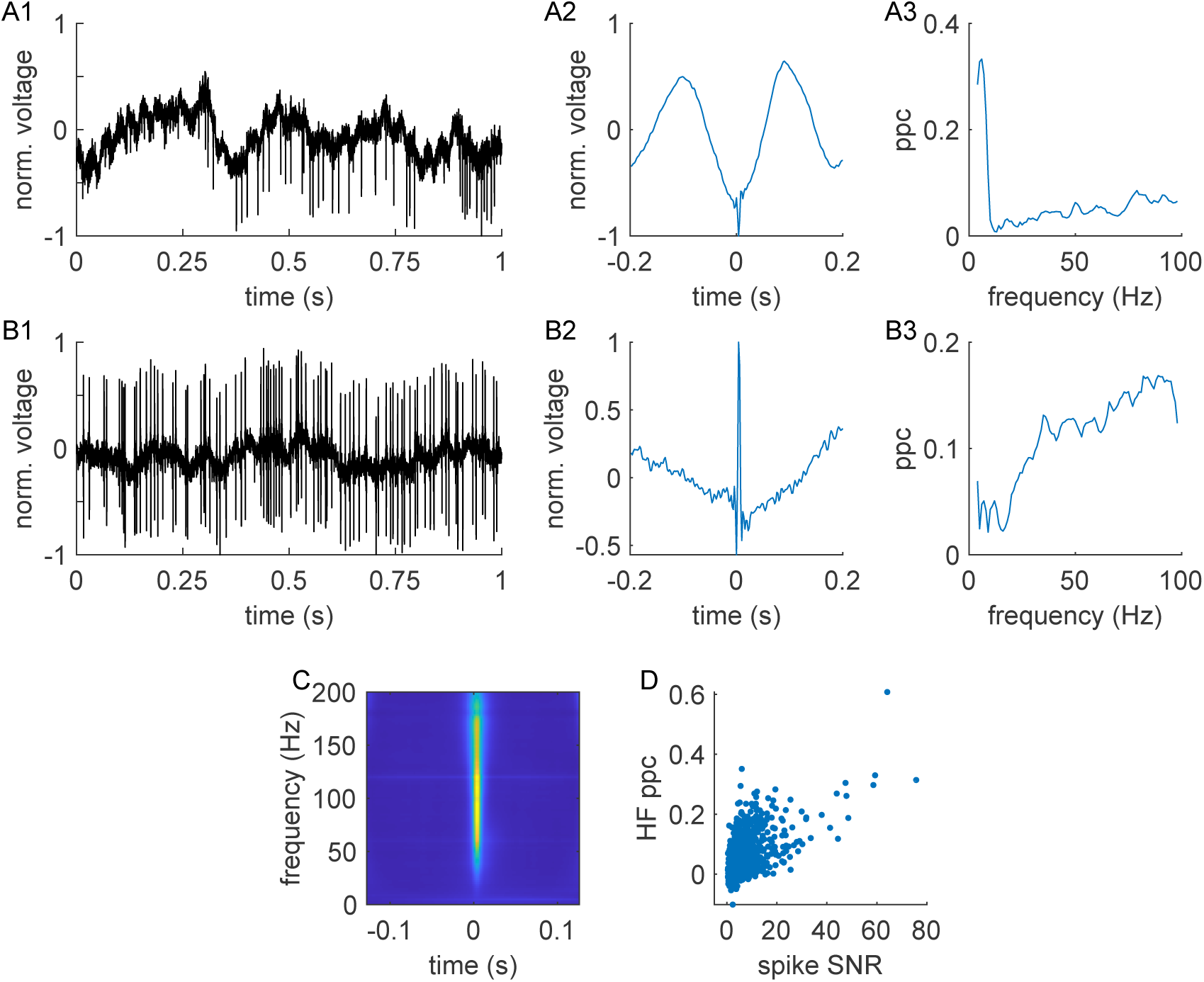
Residual spike waveforms in LFP. (A1) Example snippet of raw continuous voltage before either high-pass filtering (to extract action potentials) or low-pass filtering (to extract LFP). (A2) Spike-triggered LFP waveform (after low-pass filtering) from the example recording shown in A1. (A3) Pairwise phase consistency for the current example. (B) Additional example, same as A. (C) Average spectrogram, calculated via matching pursuit, across all LGN recordings. (D) Broadband, high-frequency spike-field phase synchrony was strongly related to the underlying signal-to-noise ratio of the LGN spike waveform.

To quantify spike-field phase synchrony and avoid the influence of the spike artifact, we applied the following reasoning. The strength of spike-field synchrony should be proportional to the underlying oscillatory power. Thus, there should be a higher degree of synchrony during periods when the oscillatory power is high compared to periods where it does not reach statistical significance. By comparison, the influence of the spike artifact should not depend upon the strength of the underlying neural oscillation. We therefore calculated spike-field ppc for periods of both strong and weak oscillatory power (Figure 5A-C, red and blue curves, respectively) and subtracted these two curves to obtain a spike artifact-free estimate of spike-field phase synchrony (Figure 5D and E). Across the sample of LGN recordings, the majority of cells had weak, statistically nonsignificant, spike-field phase synchrony, even when the analysis was restricted to epochs of strong beta-band activity (Figure 5F, see example C; ppc_strong_ = 0.001±5.3*10^−4^, ppc_weak_ = 0.0006±1.7*10^−4^). However, a minority of LGN recordings exhibited statistically significant beta-band, spike-field synchrony (Figure 5, see examples A and B), and the percentage of cells increased during epochs of strong beta-band activity (97/397, 24.4%, ppc_strong_ = 0.016±0.003) relative to epochs of weak beta-band activity (33/397, 8.3%, ppc_weak_ = 0.012±0.003, p < 10^−5^) (Figure 5E-G). Thus, if beta oscillations are indeed a feed-forward, thalamocortical signal from the LGN to V1, then the signal is likely conveyed by a minority of the LGN neurons.

**Figure 5:**
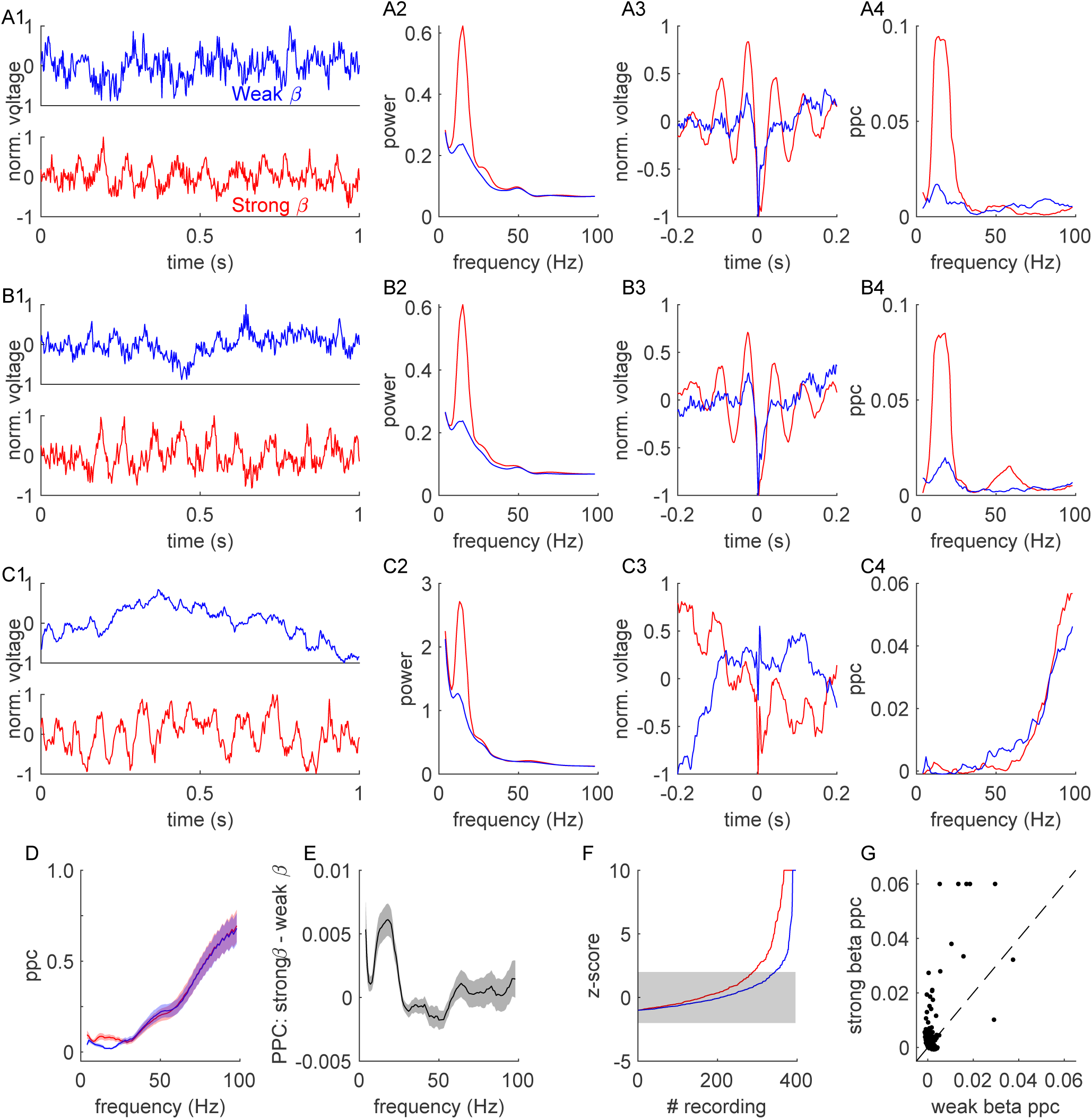
Beta-band spike-field phase synchrony in the LGN. (A1) Example snippets of weak (top trace, blue) and strong (bottom trace, red) beta band activity in LGN LFP. (A2) Example power spectra for periods of weak and strong beta band activity (blue and red traces, respectively). (A3) Example spike-triggered average LFP waveforms from same periods of weak and strong beta-band activity. (A4) Example PPC curve for the same periods of weak and strong beta-band activity. (B-C) Additional examples, same as A. (D) Average ppc curves across all LGN recordings, separated into periods of weak and strong beta-band activity. (E) Difference PPC curve: strong – weak. (F) Distribution of PPC z-scores for periods of weak and strong beta band activity. Values outside of the gray shaded region were considered significant. (G) Scatter plot of beta band PPC values for periods of weak and strong beta-band activity. For visualization purposes, values were capped at 0.06.

### Thalamocortical beta-band oscillations are suppressed by active visual processing

If beta-band thalamocortical oscillations are a mechanism that facilitates the transmission of retinal information to the cortex, then it seems reasonable to expect that both LGN oscillatory strength and LGN-V1 field-field phase synchrony would increase during periods when (1) visual stimulation is high, (2) spatial attention is directed toward the recorded retinotopic location, and (3) behavioral arousal is high. Each of these is examined below, and the results do not support the conclusion that thalamocortical beta-band oscillations are a mechanism underlying active visual processing. Instead, these oscillations are more consistent with a signature of suppression that is active when visual and behavioral processing demands are low or the demands for distractor suppression are high.

To quantify the influence of visual stimulation on thalamocortical oscillations, we first investigated the influence of stimulus strength on beta-band oscillations in the geniculocortical pathway. For this analysis, epochs of visual stimulation with sine wave gratings were divided into strong and weak classifications based on the stimulus contrast and size. Gratings that were larger than 1.5° (as large as 10°) and 50% contrast (up to 100% contrast) were considered “strong stimuli”, while gratings smaller than 1.0° (as small as 0.3°) and less than 25% contrast (as low as 5% contrast) were considered “weak stimuli”. As can be seen from both the examples and the sample average, both LGN beta-band oscillatory power (Figure 6A-E, index_ßlow_ = −0.05±0.006, p < 10^−5^; index_ßhigh_ = −0.05±0.01, p < 10^−4^) and LGN-V1 beta-band phase synchrony (Figure 7A-E; index_ßlow_ = −0.08±0.02, p < 0.001; index_ßhigh_ = −0.12±0.02, p < 10^−5^) decreased during periods of strong visual stimulation compared to periods of weak visual stimulation. Importantly, the decrease in both measures was restricted to the beta-frequency band and was not observed in other frequency bands (Figure 6F and Figure 7F). Additionally, when the analysis was restricted to recordings with significant alpha-band LGN-V1 phase synchrony (Figures 6F and 7F), there was a significant increase in alpha-band oscillatory strength (index_α_ = 0.07±0.03, p = 0.011) and LGN-V1 phase synchrony (index_α_ = 0.17±0.05, p = 0.0017) during the presentation of strong visual stimuli.

**Figure 6:**
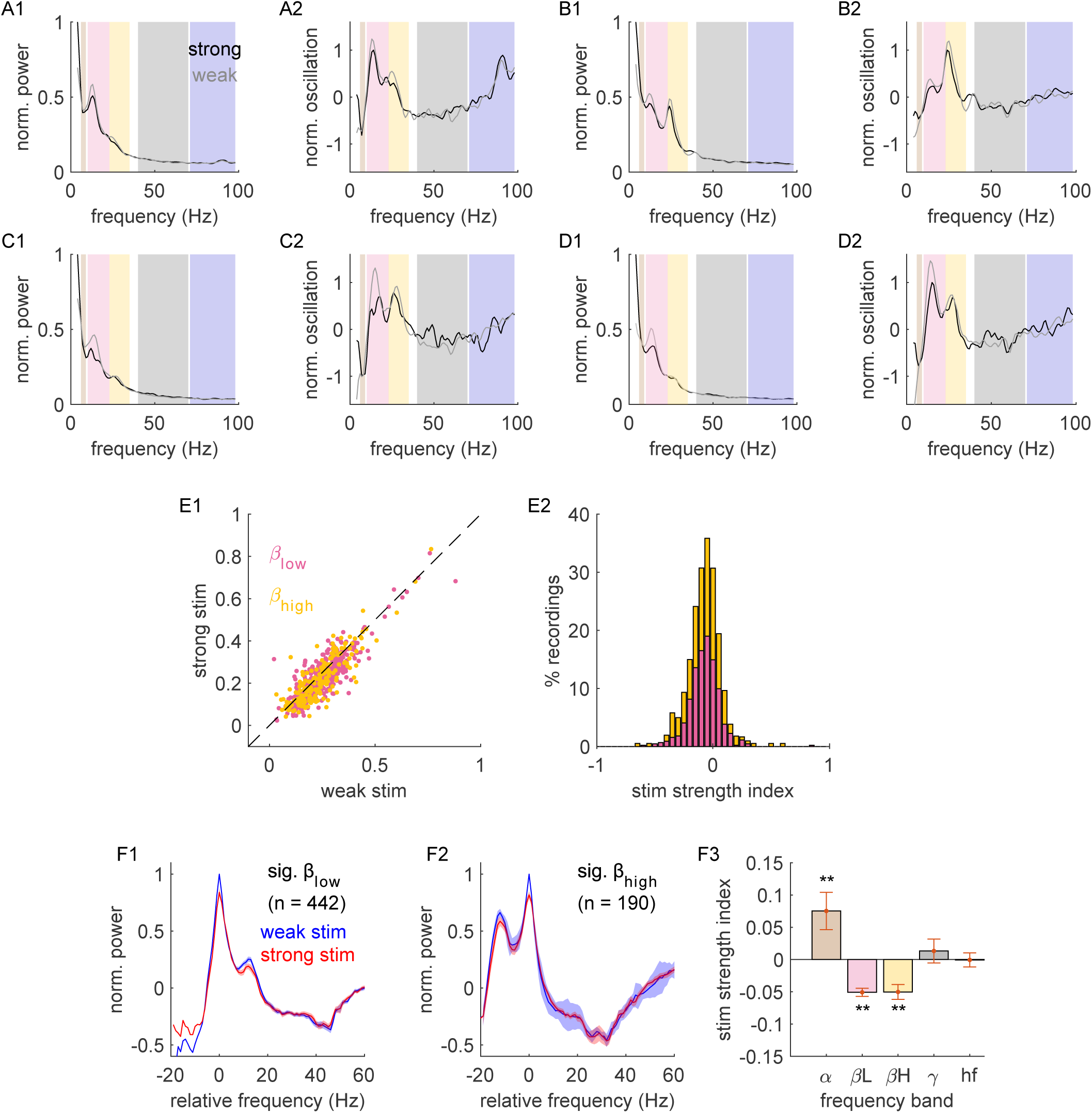
Visual stimulation suppresses beta-band oscillatory strength in the LGN (A1) Example LGN power spectra for weak and strong visual stimuli, gray trace and black trace respectively. (A2) LGN oscillatory power for current example. (B-D) additional examples, same as A. (E1) Sample distribution for LGN beta-band oscillatory power: weak vs strong visual stimulation (pink = low beta band; yellow = high beta band). (E2) Sample distribution of stimulus strength index for LGN beta-band oscillatory power (index = (strong – weak)/(strong+weak)). (F1) Average LGN oscillatory power curves (blue = weak stim, red = strong stim) for LGN recordings with significant low beta-band oscillatory strength. Frequency is defined relative to the peak low-beta band LGN oscillatory power (separately for each curve). (F2) Same as F1 for LGN recordings with significant high beta-band oscillatory strength (F3) Frequency band specific index values (** = p< 0.01).

**Figure 7:**
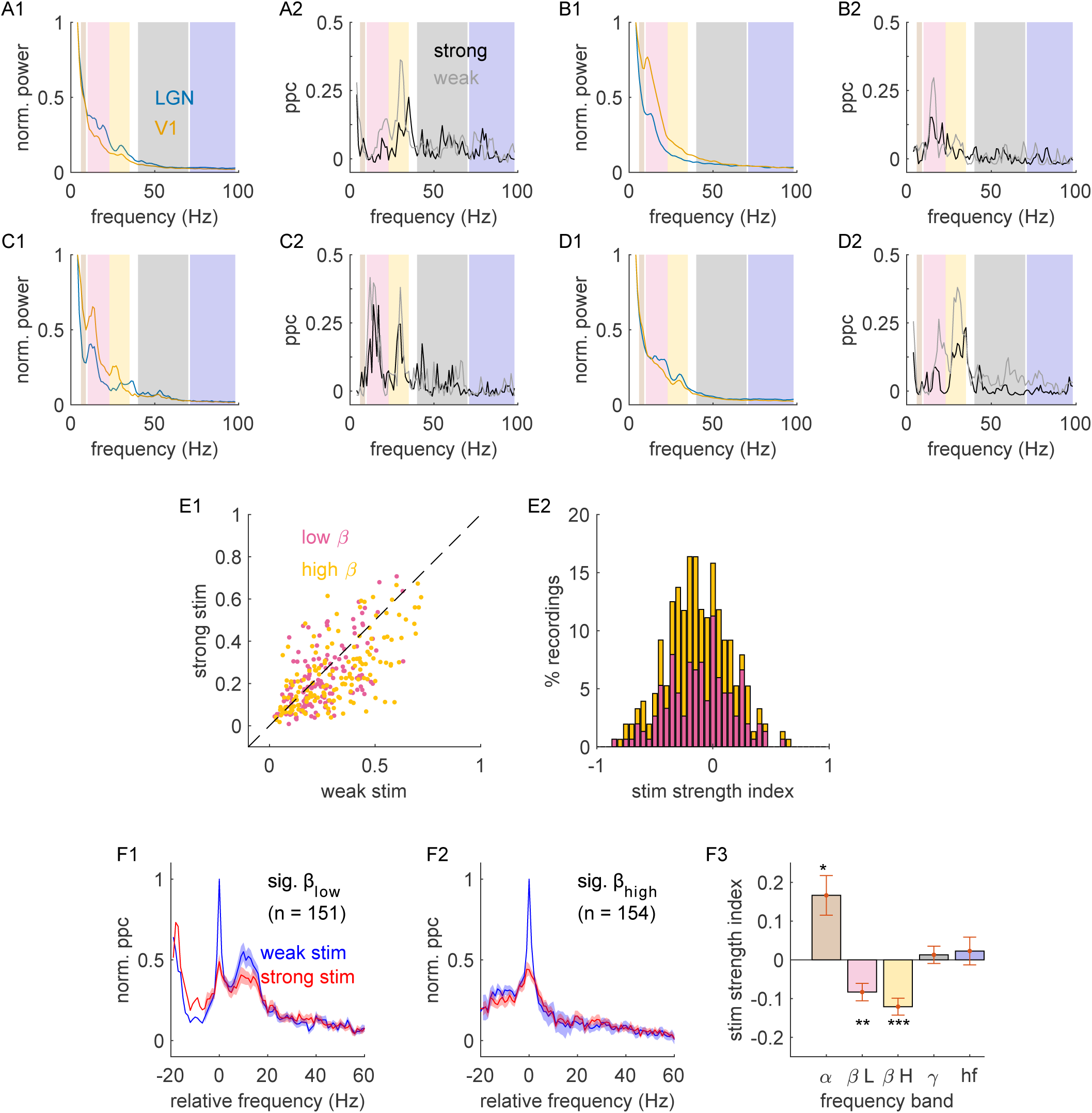
Visual stimulation suppresses beta band LGN-V1 phase synchrony. (A1) Example LGN and V1 power spectra during weak visual stimulation, blue and orange curves respectively. (A2) Example LGN-V1 field-field PPC curves: weak stimulation = gray trace, strong stimulation = black trace. (B-D) Additional examples, same as A. (E1) population distribution for LGN-V1 beta-band PPC: weak vs strong visual stimulation (pink dots = high beta-band, yellow dots = high beta-band). (E2) Population distribution of stimulus strength index for LGN-V1 beta-band PPC (index = (strong – weak)/(strong-weak)). (F1) Average LGN-V1 PPC (blue = weak stim, red = strong stim) for LGN recordings with significant low beta-band LGN-V1 phase synchrony. Frequency is defined relative to the peak low beta-band LGN oscillatory power (separately for each curve). (F2) Same as F1 for paired recordings with significant high beta-band LGN-V1 phase synchrony (F3) Frequency band specific index values (* = p<0.05, ** = p< 0.01, *** = p< 10^−5^).

Oscillatory activity throughout the brain is modulated by changes in behavioral state. This includes changes in behavioral arousal as well as changes in cognitive demands. Previous studies have demonstrated that the LGN is sensitive to levels of behavioral arousal (Livingstone and Hubel, 1981; Crombie et al., 2024). Thus, we next investigated the influence of arousal on thalamocortical beta-band oscillations (Figure 8). Towards this goal, we monitored pupil size in a subset of recordings for two monkeys as an indirect yet reliable physiological measure of overall behavioral arousal (Reimer et al., 2014; McGinley et al., 2015; Vinck et al., 2015; Crombie et al., 2024). From these measurements, we divided trials into low- and high-arousal states, representing the bottom and top 25^th^ percentiles, respectively.

**Figure 8:**
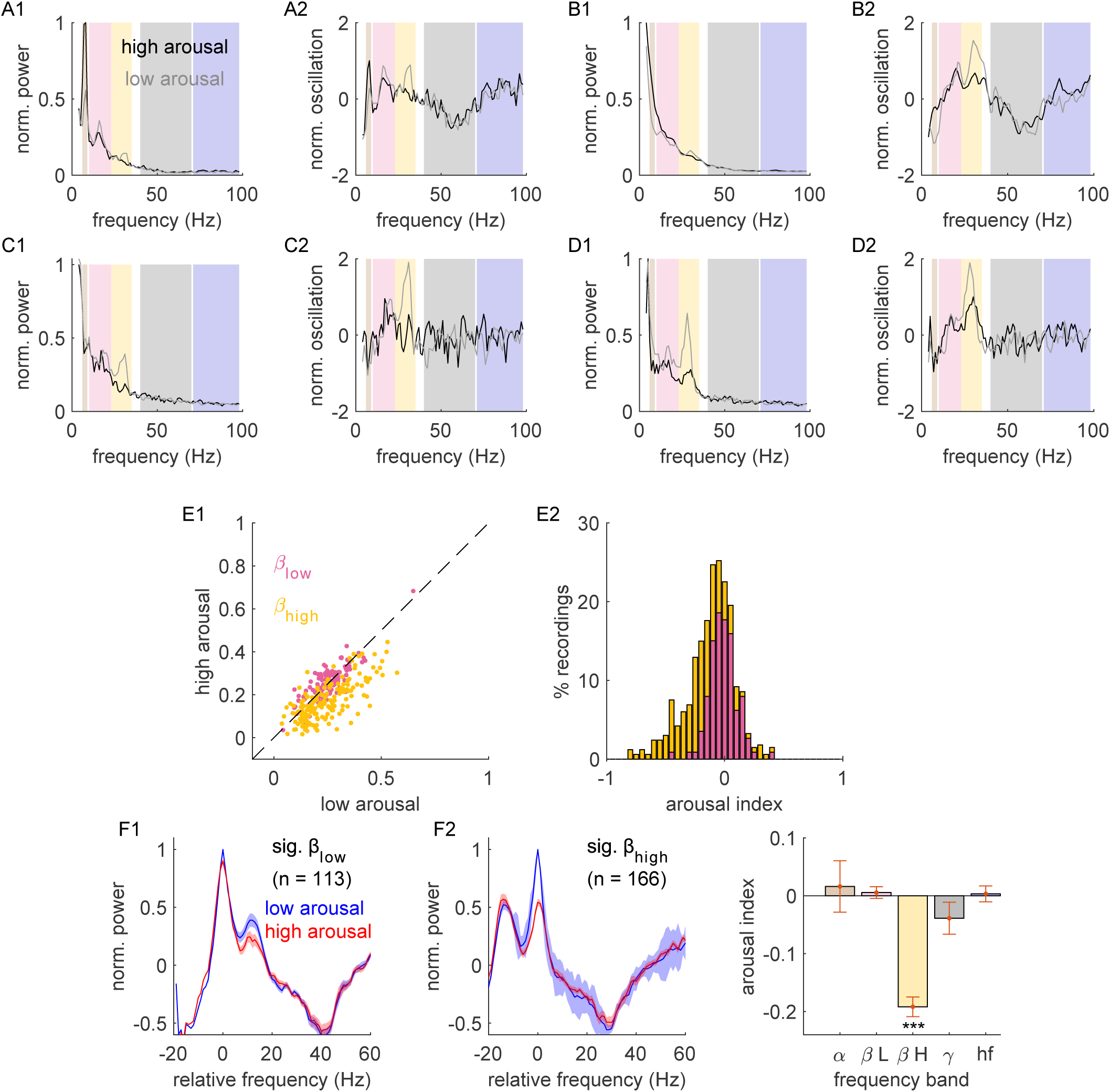
Behavioral arousal suppresses beta-band oscillations in the LGN. (A1) Example LGN power spectra for low and high arousal trials, gray trace and black trace respectively. (A2) LGN oscillatory power for current example. (B-D) additional examples, same as A. (E1) Sample distribution for LGN beta-band oscillatory power: low vs high arousal (pink = low beta band; yellow = high beta band). (E2) Sample distribution of arousal index for LGN beta-band oscillatory power (index = (high – low)/(high+low)). (F1) Average LGN oscillatory power curves (blue = low arousal, red = high arousal) for LGN recordings with significant low beta-band oscillatory strength. Frequency is defined relative to the peak low beta-band LGN oscillatory power (separately for each curve). (F2) Same as F1 for LGN recordings with significant high beta-band oscillatory strength (F3) Frequency band specific index values (*** = p< 10^−5^).

Increases in behavioral arousal, as inferred through increases in pupil size, were associated with decreases in thalamocortical oscillations. We found a decrease in the high beta-band LGN oscillatory power (Figure 8A-E; index_ßhigh_ = −0.19±0.02, p < 10^−5^ index_ßlow_ = 0.005±0.01, p = 0.58) as well as a decrease in the low and high beta-band, LGN-V1phase synchrony (Figure 9A-E; index_ßlow_ = −0.12±0.04, p = 0.001; index_ßhigh_ = −0.13±0.02, p < 10^−5^). Importantly, these decreases were specific to the beta-frequency band and were not representative of an overall decrease in network activity (Figure 8F and Figure 9F).

**Figure 9:**
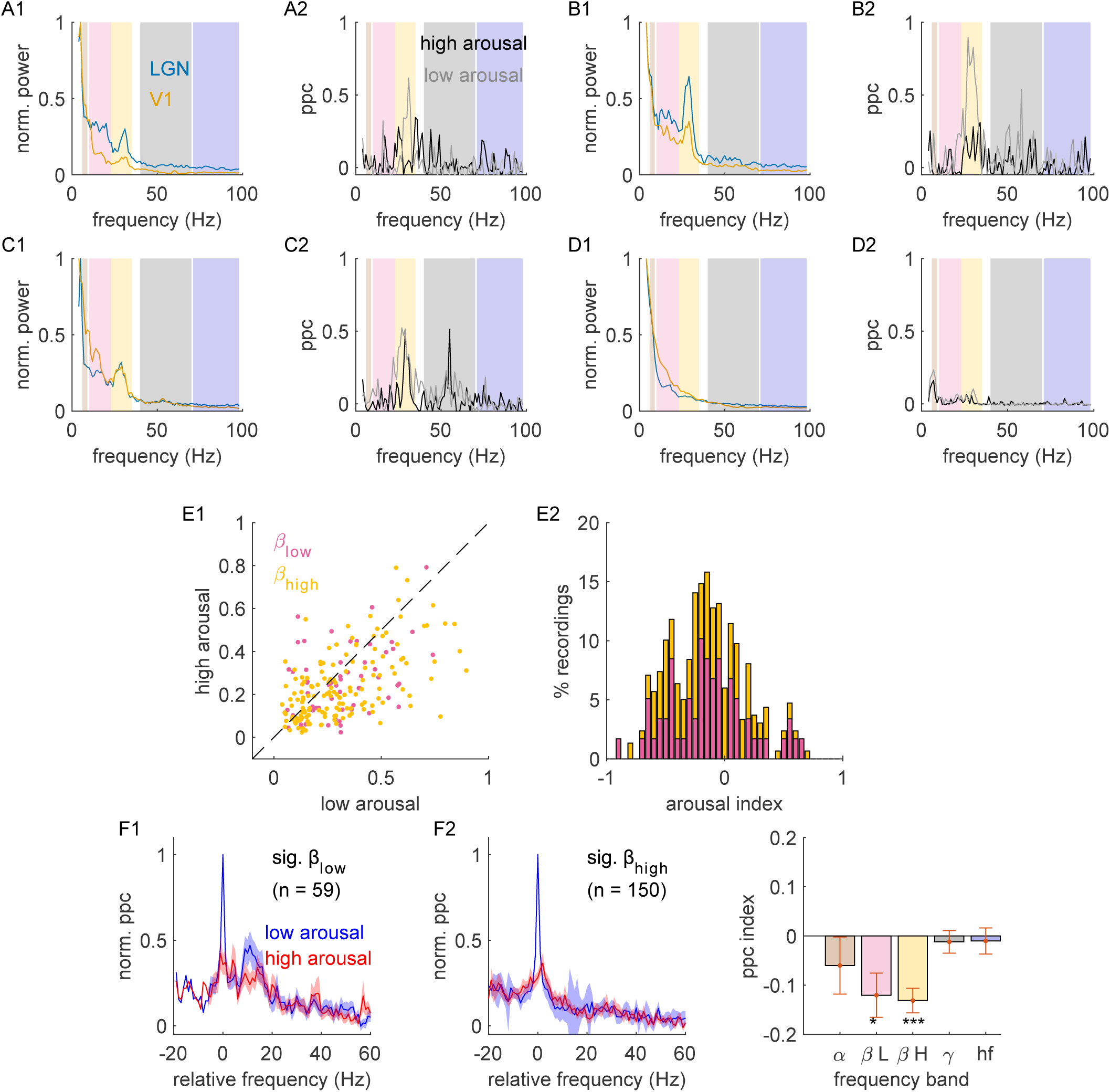
Behavioral arousal suppresses beta band LGN-V1 phase synchrony. (A1) Example LGN and V1 power spectra during low behavioral arousal, blue and orange curves respectively. (A2) Example LGN-V1 field-field PPC curves: low arousal = gray trace, high arousal = black trace. (B-D) Additional examples, same as A. (E1) population distribution for LGN-V1 beta band PPC: low vs high behavioral arousal (pink dots = high beta-band, yellow dots = high beta-band). (E2) Population distribution of arousal index for LGN-V1 beta-band PPC (index = (high – low)/(high-low)). (F1) Average LGN-V1 PPC (blue = low arousal, red = high arousal) for LGN recordings with significant low beta-band LGN-V1 phase synchrony. Frequency is defined relative to the peak low beta=band LGN oscillatory power (separately for each curve). (F2) Same as F1 for paired recordings with significant high beta-band LGN-V1 phase synchrony (F3) Frequency band specific index values (* = p<0.05, ** = p< 0.01, *** = p< 10^−5^).

Finally, we recorded from two macaque monkeys while they performed a spatial attention task where their attention was either directed toward or away from the retinotopic position of the LGN receptive field (see Materials and Methods). Importantly, the animals displayed psychophysical behavioral signatures of performing the task as anticipated. Specifically, they were faster and more accurate at reporting changes at the attended location (valid cue trials) compared to changes at the unattended location (invalid cue trials) (Alitto et al., 2025). For these experiments, recordings were only made from the LGN, so results are limited to the influence of spatial attention on LGN oscillatory strength.

Shifting spatial attention toward the retinotopic position of the LGN receptive field caused a small, but significant, decrease in thalamocortical oscillations as assessed via LGN beta-band oscillatory strength (Figure 10; index_ßlow_ = −0.02±0.006, p = 0.002; index_ßhigh_ = −0.04±0.02, p = 0.007). Importantly, the changes to LGN oscillatory power were specific to the beta-frequency band and did not represent an overall decrease in network activity (Figure 10F3). In contrast, we saw a small but significant increase in the firing rate of LGN neurons when attention was shifted toward the LGN receptive field, relative to when attention was shifted away from the LGN receptive field (Alitto et al., 2025). We speculate that the difference in firing rate may be associated with the change in beta-band oscillatory power, which may serve to suppress the transmission of retinal signals that encode visual distractors, such as the unattended drifting grating.

**Figure 10:**
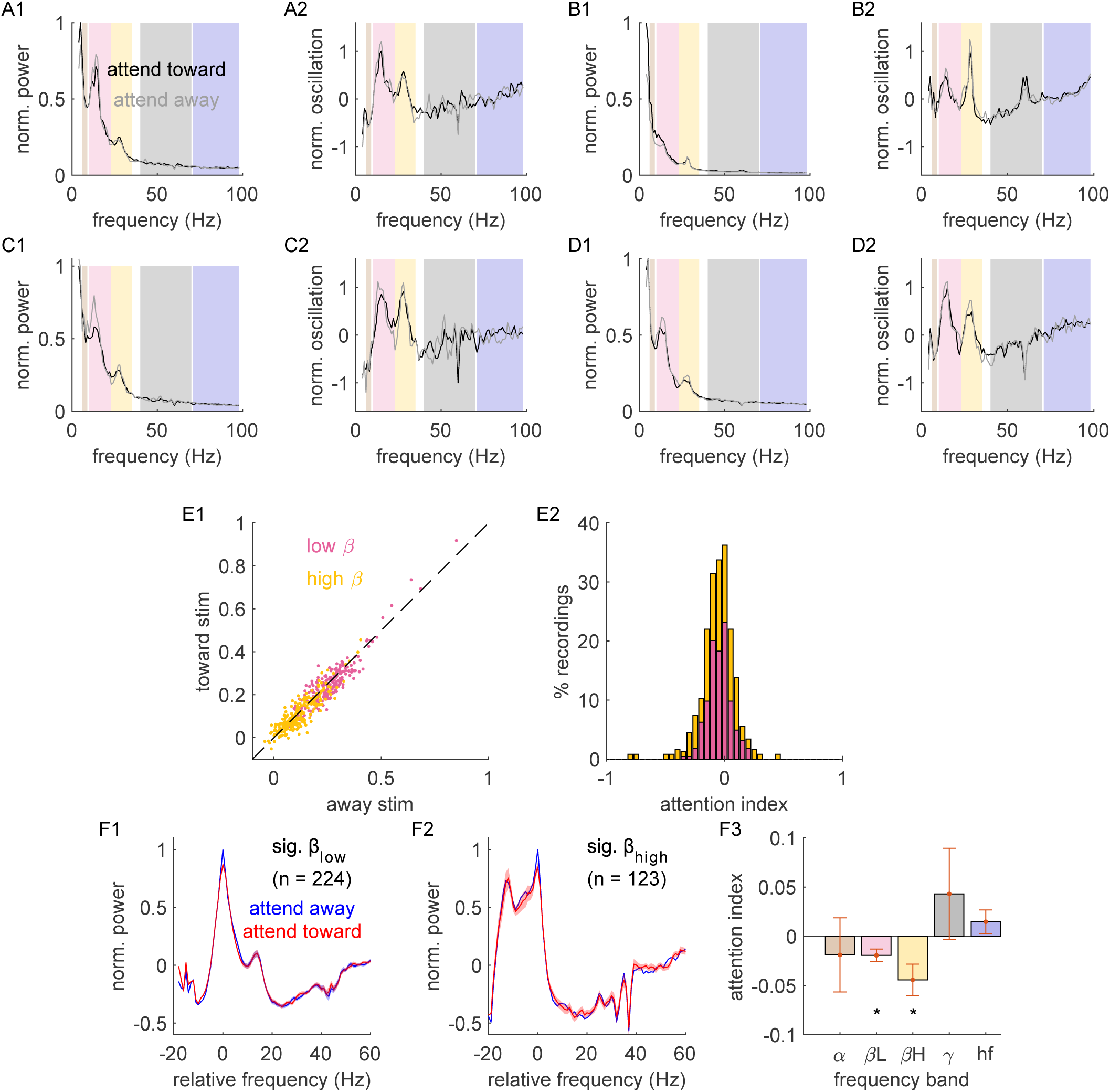
Shifts in spatial attention modulate beta-band activity in the LGN. (A1) Example LGN power spectra when spatial attention was directed away from and toward an LGN receptive field, gray trace and black trace, respectively. (A2) Example LGN oscillatory power. (B-D) additional examples, same as A. (E1) Sample distribution for LGN beta-band oscillatory power: away vs toward the LGN receptive field power (pink = low beta band; yellow = high beta band) (E2) Population distribution of spatial attention index for LGN beta-band oscillatory power (index = (toward – away)/(toward+high)). (F1) Average LGN oscillatory power curves (blue = away, red = toward) for LGN recordings with significant low beta-band oscillatory strength. Frequency is defined relative to the peak low beta-band LGN oscillatory power (separately for each curve). (F2) Same as F1 for LGN recordings with significant high beta-band oscillatory strength (F3) Frequency band specific index values (* = p< 0.05).

In summary, this study revealed prominent beta-band oscillations in the LGN that are coherent with V1 and are statistically consistent with a feedforward, thalamocortical process from the LGN to V1. These oscillations are suppressed by visual stimulation, behavioral arousal, and shifts in spatial attention toward the LGN receptive field, indicating they do not represent a mechanism underlying active visual processing.

## DISCUSSION

Consistent with Bastos et al. (2014), we observed robust beta-band oscillations in the lateral geniculate nucleus (LGN) that were coherent with activity in primary visual cortex (V1) in a manner that was statistically consistent with a feedforward thalamocortical signal. These oscillations influenced spike timing in a subset of LGN neurons, suggesting a plausible mechanism for transmitting rhythmic activity to the cortex. Critically, however, we found that LGN-V1 beta-band oscillations were suppressed, rather than enhanced, during periods of increased network engagement. Specifically, this suppression occurred during strong visual stimulation, elevated arousal, and covert shifts of spatial attention toward the LGN receptive field. This inverse relationship between beta-band power and sensory-cognitive demands strongly suggests that thalamocortical oscillations do not enhance information transmission between the LGN and V1. Instead, our findings are more consistent with the robust literature on cortical alpha-band rhythms, which are understood to reflect active suppression of task-irrelevant input and are modulated by attention, arousal, and perceptual load. In the sections below, we expand on this proposed functional analogy between these thalamocortical beta oscillations and cortical alpha oscillations, discuss their shared sensitivity to behavioral context, and explore potential circuit mechanisms that may support this dynamic form of suppression.

### Oscillatory mechanisms of signal suppression in the visual system

EEG and LFP recordings from the awake brain, in both humans and animals, are often dominated by cortical alpha-band oscillations (Lopes da Silva et al., 1973; Başar et al., 2001; Palva and Palva, 2007; Rihs et al., 2007; Bollimunta et al., 2011; Mo et al., 2011; Haegens et al., 2015). One of the earliest observations was that alpha power increases when subjects close their eyes during quiet wakefulness, a state long associated with reduced sensory input and diminished visual processing (reviewed in Shomer, 2017). While alpha-band oscillations were once viewed as a signature of cortical idling, they are now widely believed to represent an active inhibitory mechanism that limits interference from task-irrelevant inputs and modulates cortical excitability in accordance with behavioral demands (Kelly et al., 2006; Klimesch et al., 2007; Rihs et al., 2007). This suppressive function has been observed across a range of experimental paradigms. Alpha-band power increases over task-irrelevant visual regions during spatial attention tasks (Gutteling et al., 2022; Yang et al., 2024), across unattended sensory modalities during modality-specific attention (Foxe et al., 1998), and in motor cortex during the inhibition of learned responses (Hummel et al., 2002). In working memory tasks, elevated alpha-band power suppresses interference from new visual input during the maintenance phase (Herrmann et al., 2004; Tuladhar et al., 2007; Jensen and Mazaheri, 2010). Importantly, alpha-band activity is not merely present during suppression, but actively tracks the demands of suppression. For example, in spatial attention tasks, alpha-band power increases more strongly over regions representing distractors when those distractors are physically similar to targets (Gutteling et al., 2022). This observation aligns with perceptual load theory, which posits that the degree of distractor suppression increases with task difficulty. It is also now well established that alpha-band activity influences perceptual awareness. Trial-to-trial variability in alpha-band power predicts whether near-threshold visual stimuli are consciously detected (Di Gregorio et al., 2022).

Based on the modulation of beta-band oscillations in our dataset by both arousal and spatial attention, we propose that these oscillations in the macaque geniculocortical pathway are functionally analogous to the alpha-band oscillations commonly observed in human cortical EEG. Similar to cortical alpha, LGN-V1 beta-band oscillations were strongest during states of low engagement and were reduced when sensory or attentional demands increased. This pattern was evident across multiple behavioral contexts. First, during passive fixation and the presentation of behaviorally irrelevant visual stimuli, beta-band power increased during periods of low arousal, as indexed by pupil size. While fixation on a 1° target requires minimal but sustained engagement, we interpret these oscillations as consistent with quiet wakefulness or partial disengagement. Second, during a spatial attention task that required significantly more engagement, we observed that beta-band oscillations were stronger when the distractor, rather than the target, overlapped the LGN receptive field. This spatial selectivity closely mirrors the retinotopically specific increases in alpha-band power observed in visual cortex when attention is directed away from a given location. Together, these findings suggest that LGN-V1 beta-band oscillations serve as a dynamic index of functional inhibition, suppressing inputs that are behaviorally irrelevant or unattended. We emphasize that our data demonstrate a correlational relationship consistent with this suppressive function. Future experiments involving causal perturbation will be necessary to establish whether beta-band oscillations in the LGN actively gate information flow to the cortex and/or modulate perceptual sensitivity.

Although we conclude that the LGN-V1 beta-band oscillations observed in macaques are functionally analogous to cortical alpha-band oscillations, we refrain from labeling them as “alpha” due to frequency range considerations. Specifically, these oscillations often extended above 30 Hz, and there is no precedent in the literature for referring to such high-frequency activity as alpha. To maintain consistency with existing nomenclature, we instead adopt a conventional labeling scheme: low-beta (11–24 Hz) and high-beta (25–35 Hz) (HajiHosseini et al., 2012; Zhang et al., 2019). An alternative approach would have been to label the low-frequency component as alpha and the high-frequency component as beta, as has been done in some prior studies (e.g., Schmiedt et al., 2014). However, this binary distinction would imply a functional dissociation between the two sub-bands—an interpretation not supported by our data.

Importantly, while the oscillations we report occupy the beta-frequency range, their peak frequencies varied across animals and sometimes overlapped with traditional alpha-band frequencies (e.g., 10–14 Hz). This observation is consistent with prior findings showing substantial inter-individual variability in alpha frequency (Benwell et al., 2019) and even trial-by-trial variability within individuals (Minami et al., 2020). Moreover, recent work suggests that functionally analogous rhythms may appear at lower frequencies in other species, such as 3–5 Hz rhythms in rodent visual thalamus (Nestvogel and McCormick, 2022). Taken together, these findings highlight the view that the functional role of an oscillation cannot be fully determined by its frequency alone. Ultimately, while frequency-based labels offer a useful shorthand, they are only meaningful to the extent that they reflect shared mechanisms and functions across experimental contexts.

### The neural origin of LGN-V1 oscillations

The LGN and V1 are interconnected not only via feedforward excitatory projections from the LGN to V1 but also through corticothalamic feedback projections originating from layer 6 of V1. This feedback provides both direct excitation to LGN relay neurons and indirect inhibition via the thalamic reticular nucleus (TRN) and local interneurons(Sherman and Guillery, 2006). Given this bidirectional architecture, distinguishing whether oscillations observed in the LGN-V1 circuit arise through feedforward or feedback (or both) processes is nontrivial.

To probe directionality, we applied a phase slope index (psi, which revealed that LGN-V1 beta-band synchrony was statistically consistent with a feedforward signal from LGN to V1. While psi is less susceptible to spurious inferences than other metrics like Granger causality (Nolte et al., 2008), it is ultimately a statistical inference based on relative phase delays and cannot alone establish the origin of the observed oscillations. Indeed, evidence from the broader literature suggests that alpha-band oscillations—functionally similar to the LGN-V1 beta-band oscillations described here—can arise from both thalamic and cortical sources. In thalamus, rhythmic inhibition from the TRN and high-threshold burst firing in specialized relay neurons have been implicated in alpha-frequency generation (Hughes and Crunelli, 2005). In cortex, infragranular activity, particularly in cortical layer 5, has been proposed as a source of alpha-band oscillatory activity, especially during attentionally relevant processing (Bollimunta et al., 2008; Spaak et al., 2012).

We therefore propose that LGN-V1 beta-band oscillations arise from both thalamic and cortical generators, which are then synchronized via recurrent thalamocortical loops. Such hybrid origins have been described for alpha-band rhythms and may also account for the behavioral sensitivity and feedforward dynamics of the beta-band oscillations found in this study.

## Acknowledgments

We thank K.E. Neverkovec, Jeffrey S. Johnson, D.J. Sperka, and R. Oates for expert technical assistance. We also thank Rober E. Boshra and Nancy Kopell for their comments on the manuscript.

