## Supplementary figures and images for "Dynamic Modulation of Beta-Band Oscillations in the LGN and Their Role in Visual Processing"

### Supplemental Figure S1

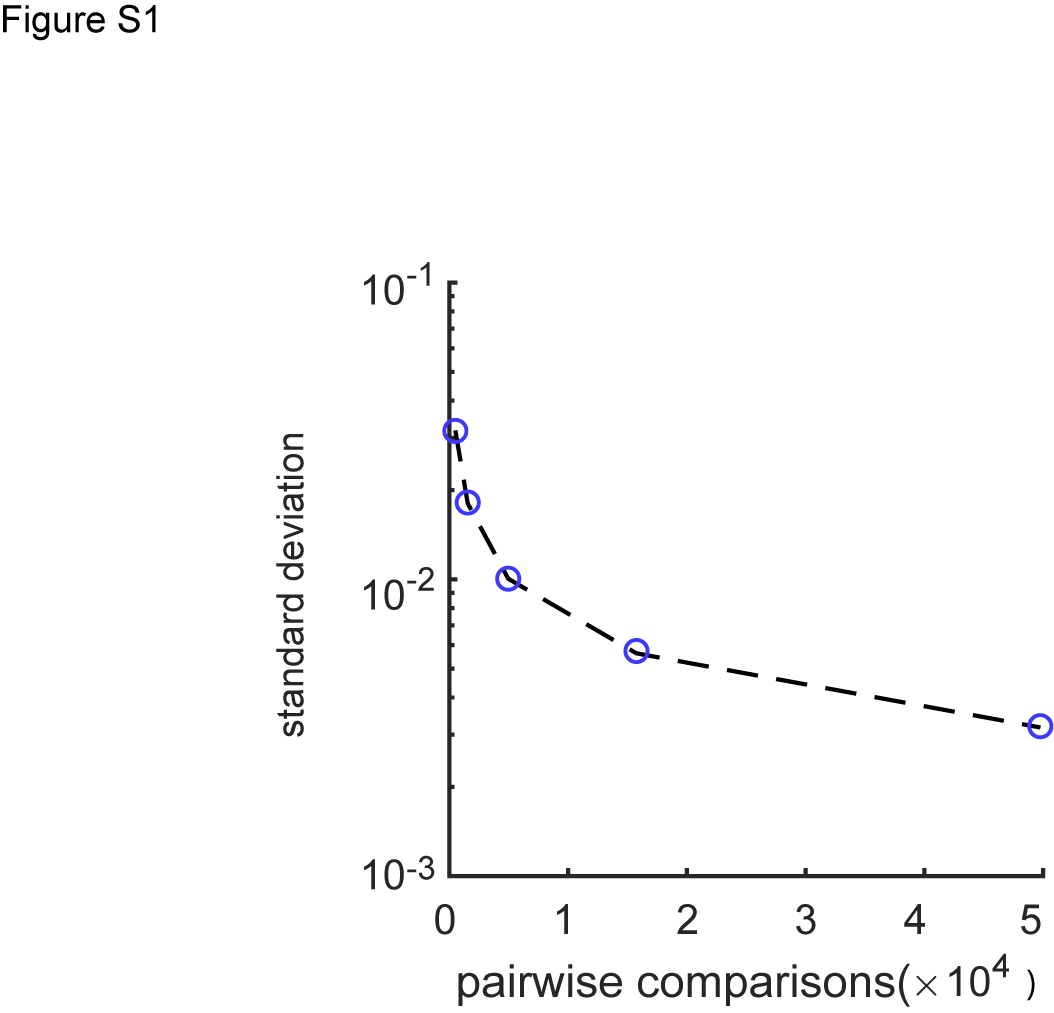
