## Supplemental Figure Legend for "Dynamic Modulation of Beta-Band Oscillations in the LGN and Their Role in Visual Processing"

Figure S1: PPC standard deviation. As derived in the methods, we defined the standard deviation of the ppc null hypothesis (i.e., a random distribution of phases) as $\frac{1}{\sqrt{2N}}$ where N = number of pairwise comparisons. To verify the accuracy of this calculation we simulated ppc given the null hypothesis across a range of Ns. The standard deviation calculated directly from these simulations (5000 repeats each, blue circles) matches very closely with the analytically defined values (dashed line).
